# Altered mechanical properties of astrocytes lacking MLC1; implications for the leukodystrophy MLC

**DOI:** 10.1101/2025.02.19.639082

**Authors:** Quinty Bisseling, Emma M.J. Passchier, Nelda Antonovaite, Serena Camerini, Maria S. Brignone, Amélie Freal, Huibert D. Mansvelder, Elena Ambrosini, Marjo S. van der Knaap, Rogier Min

## Abstract

Loss of function of the astrocyte protein MLC1 causes Megalencephalic Leukoencephalopathy with subcortical Cysts (MLC), a leukodystrophy characterized by white matter edema and slow neurological deterioration. MLC1 dysfunction leads to swelling of perivascular astrocyte endfeet and an impaired attachment of endfeet to blood vessels. In isolated primary astrocytes, loss of MLC1 hinders recovery of astrocytes from cell swelling, but the cellular function of MLC1 is not completely understood. MLC1 modulates gating of mechanosensitive ion channels involved in volume regulation. The cytoskeleton plays a crucial role in cell volume regulation, and interactions between the cytoskeleton and cell membrane affect the properties of mechanosensitive ion channels. Therefore, we investigated whether primary *Mlc1*-null mouse astrocytes show a disruption in their mechanical properties. We measured mechanical properties of cultured primary astrocytes with an indentation technique and demonstrated that *Mlc1*-null astrocytes are softer than wild-type astrocytes. Proteomic analysis confirmed dysregulation of several cytoskeleton-related pathways in *Mlc1*-null astrocytes. Confocal imaging revealed that organization of the actin cytoskeleton is unaffected. Instead, we observed alterations in focal adhesions, which aid in relaying mechanical forces between the cytoskeleton, cell membrane, and the extracellular matrix (ECM). Together, our findings reveal that the mechanical properties of *Mlc1*-null astrocytes are altered, and that disrupted cytoskeleton-membrane-ECM interactions potentially play a role in the disease. Modulators of astrocyte mechanobiology might therefore hold promise for MLC therapy development.

## 1 | INTRODUCTION

Astrocytes play a crucial role in the regulation of fluid flow in the brain. They form specialized compartments called endfeet, which surround the entire brain microvasculature (Mathiisen et al., 2010). Endfeet are equipped with channels, pumps and membrane proteins essential for brain ion and water homeostasis. Dysfunction of endfeet leads to neurological diseases (Min & van der Knaap, 2018), such as the leukodystrophy Megalencephalic Leukoencephalopathy with subcortical Cysts (MLC, OMIM 604004). MLC is characterized by disturbed brain ion and water homeostasis (Ridder et al., 2011; van der Knaap et al., 2012). Patients have macrocephaly due to chronic brain white matter edema, and they have epilepsy, mild cognitive disability and progressive motor impairment (van der Knaap et al., 1995; Singhal et al., 1996). In MLC, astrocyte endfeet are swollen and fluid-filled vacuoles can be found in myelin and, to a lesser degree, in astrocyte endfeet (van der Knaap et al., 2012; Dubey et al., 2015). In most MLC patients the disease is caused by a loss of function of the membrane protein MLC1 (Leegwater et al., 2001), which is highly expressed in astrocyte endfeet (Boor et al., 2005). Defects in several other endfoot proteins that interact with MLC1 also cause MLC, specifically the glial cell adhesion molecule GlialCAM, the orphan G protein-coupled receptor GPRC5B and the main water channel of the brain, Aquaporin-4 (AQP4) (López-Hernández, Ridder et al., 2011; Passchier et al., 2023). All four MLC-related proteins have been linked to astrocyte volume regulation. Primary *Mlc1*-null mouse astrocytes as well as patient lymphoblasts show a defect in regulatory volume decrease (RVD), the process that allows cells to recover from osmotic swelling. This defect has been attributed to dysfunction of mechanosensitive ion channels such as the transient receptor potential cation channel subfamily V member 4 (TRPV4) and the volume-regulated anion channel (VRAC) (Ridder et al., 2011; Lanciotti et al., 2012; Dubey et al., 2015; Jentsch, 2016; Passchier et al., 2023). The exact function of MLC1 and the mechanisms by which MLC1 modulates ion channel activity and astrocyte volume regulation remain unclear (Elorza-Vidal et al., 2018).

Volume regulation relies on cellular mechanical properties, which are determined by the cytoskeleton and its interactions with the membrane and the extracellular matrix (ECM). The cytoskeleton consists of intermediate filaments, microtubules and actin. Actin filaments can interact with macromolecular complexes, like focal adhesions (FAs), providing a direct link between the intracellular cytoskeleton and the ECM. The cytoskeleton plays an important role in astrocyte volume regulation through modulation of mechanosensitive volume-regulated ion channels (Levitan et al., 1995; Lascola & Kraig, 1996; Lascola et al., 1998; Turovsky et al., 2020). Several studies suggest a role for MLC-related proteins in modulating cytoskeleton-membrane-ECM interactions, thereby affecting cellular mechanical properties. MLC1 and GlialCAM are predominantly present at astrocyte-astrocyte junctions, where cis and trans interactions between GlialCAM proteins support cell-to-cell adhesion (Teijido et al., 2007; López-Hernández, Ridder et al., 2011; López-Hernández, Sirisi et al., 2011). MLC1 colocalizes with the actin cytoskeleton (Duarri et al., 2011), and is associated with the dystrophin-associated glycoprotein complex (DAGC), which supports adhesion of perivascular endfeet to the ECM (Boor et al., 2007; Ambrosini et al., 2008). *Mlc1*-null mouse tissue shows a decrease in perivascular endfeet coverage, with endfeet not being adequately attached to the vasculature (Gilbert et al., 2021). Both MLC1 and AQP4 interact with TRPV4, which is directly connected to the actin cytoskeleton (Benfenati et al., 2011; Lanciotti et al., 2012; Jo et al., 2015). MLC1 negatively regulates actin branching by binding the Arp2/3 complex, which regulates actin nucleation (Hwang et al., 2019). Besides directly interacting with MLC1, the Arp2/3 complex transiently binds vinculin, which promotes FA formation (DeMali et al., 2002; Chorev et al., 2014). Vinculin is a large scaffolding protein that resides in the force-transduction layer of FAs and links the ECM to the actin cytoskeleton (Kanchanawong et al., 2010). Thus, multiple studies indicate interactions between MLC1 and different components that are important in regulating mechanical properties of cells. How loss of MLC1 function affects astrocyte mechanobiology has not been studied. The aim of this study is to investigate the mechanical properties of cultured primary astrocytes isolated from an MLC mouse model. We hypothesize that loss of function of MLC1 alters the structural integrity of astrocytes by disturbing cytoskeleton-membrane-ECM interactions.

## 2 | MATERIALS AND METHODS

### 2.1 | Animals

Primary astrocyte cell cultures were obtained from wild-type and transgenic *Mlc1*-null mice. All mice had a C57Bl/6J background. Generation of *Mlc1*-null mice is described in (Dubey et al., 2015). Wild-type and *Mlc1*-null mice were obtained by homozygous breeding of *Mlc1*^wt/wt^ x *Mlc1*^wt/wt^ or *Mlc1*^null/null^ x *Mlc1*^null/null^ mice, respectively. Breeding pairs were regularly refreshed from heterozygous breeding to limit drift in genetic background. Experimental procedures involving mice were in strict compliance with animal welfare policies of the Dutch government and were approved by the Institutional Animal Care and Use Committee of the Amsterdam University Medical Center, location AMC, Amsterdam, or of the Vrije Universiteit Amsterdam, depending on the location of experiments.

### 2.2 | Isolation of primary astrocytes from mice

Cortical primary astrocytes were isolated from neonatal wild-type and *Mlc1*-null mice on postnatal day 6 to day 9 (p6-p9) (protocol adapted from (McCarthy & de Vellis, 1980). Brains were isolated from the skull and cerebral cortices were dissected by removing the olfactory bulb, cerebellum, midbrain and meninges. This was done in ice-cold, sterile Hanks’ Balanced Salt Solution without calcium and magnesium, with Phenol Red (HBSS^-/-^; Gibco, 14170-088) supplemented with 1% Penicillin Streptomycin (Pen Strep; Gibco, 15140-122). Obtained neocortical tissue was minced with a sterile scalpel and incubated in TrypLE™ Express (1X; Gibco, 12605-010) supplemented with DNase I (40 µg/mL; Roche, 11284932001) in a rotator device for 25 min at 37 °C. The cell suspension was centrifuged (475 x *g*, 3 min) at room temperature (RT) and the supernatant was removed. The pellet was resuspended in complete primary astrocyte medium (DMEM/F-12 (1:1) with GlutaMAX^TM^-I and Phenol Red (Gibco, 31331-028), 10% Fetal Bovine Serum (FBS; Gibco, 10270-106), 1% Sodium Pyruvate (Gibco, 11360-039, 100 mM) and 1% Pen Strep) and centrifuged again (475 x *g*, 3 min, RT). For mechanical dissociation the cell pellet was titrated in complete primary astrocyte medium using a 5 mL and 2 mL pipette, respectively. In between and after the titration steps the cell suspension was centrifuged (475 x *g*, 3 min, RT) and resuspended in complete primary astrocyte medium. The cell suspension was filtered through a 70 µm Nylon Cell Strainer (Corning, 431751) and centrifuged (475 x *g*, 5 min, RT). Finally, the cell pellet was resuspended in complete primary astrocyte medium and transferred into poly-L-lysine (PLL)-coated (0.1 mg/mL for 1-2 h at 37 °C; Sigma, P2636) T75 flasks and incubated in a humidified CO_2_ incubator (37 °C / 5% CO_2_). Until >80% confluency was reached (∼1 week), cells were washed with Dulbecco’s Phosphate Buffered Saline (DPBS; Gibco, 14190-094) and medium was changed every 2-3 days. At >80% confluency, contaminating oligodendrocytes, microglia and precursor cells were dislodged overnight (ON) by orbital shaking (180 RPM; VWR, 89032-088). The next day, flasks were vigorously shaken by hand for 30 sec and washed in DPBS 3 times. Complete primary astrocyte medium was added, and cells were used in experiments or maintained for a week maximum in a humidified CO_2_ incubator (37 °C / 5% CO_2_) with medium changes every 3 days.

### 2.3 | Cell culture and treatments

Primary astrocytes were washed with DPBS and incubated in TrypLE™ for 5 min at 37 °C. The flask was tapped furiously by hand, cells were resuspended in complete primary astrocyte medium and used for plating or preparation of cell pellets.

*For indentation experiments*, primary astrocytes were seeded in uncoated 24-well plates (Biofil, TCP-011-024) at a density of 15.000 cells per well and used for experiments between 24 h and 48 h after plating. Before the start of the experiment cells were treated with cytochalasin D (2 µM, 5 µM or 10 µM; Sigma-Aldrich, C2618, 5 mg/mL in DMSO) diluted in complete primary astrocyte medium for 1 h at 37 °C. After treatment medium was replaced with 1 mL isotonic solution in which experiments were performed. Isotonic solution contained (in mM): 140 NaCl, 4 KCl, 2 MgCl_2_, 2 CaCl_2_, 10 HEPES, 5 D(+)-glucose (pH adjusted to 7.4 with NaOH and osmolality adjusted to 300 mOsm/kg).

*For the preparation of cell pellets for mass spectrometry*, primary astrocytes were collected into a 50 mL tube after resuspension in complete primary astrocyte medium. DPBS was added to the cell suspension to fill up the tube to 50 mL. The cell suspension was washed with DPBS 3 times by spinning down the cells to form a cell pellet (400 x *g*, 5 min, RT), then removing the supernatant and adding DPBS after each round. The cell pellets were resuspended in 10 mL DPBS, and cells were counted. After spinning down again the cell pellet was resuspended and transferred to freezer vials (VWR, 479-1256) at 2*10^6^ cells/ml and spun down (400 x *g*, 5 min, RT). Excess DPBS was removed, and the cell pellets were frozen at −20 °C and after 24 h stored at −80 °C. Cell pellets were shipped on dry ice. Each cell pellet represents one mouse brain.

*For immunofluorescence staining,* primary astrocytes were seeded on PLL-coated (0.1 mg/mL for 1-2 h at 37 °C) glass coverslips (13 mm, VWR, 631-1578) in a 24-well plate at 15.000 cells per well. 48 h after plating the cells were washed 3 times with DPBS, fixed in DPBS with 2% paraformaldehyde (PFA; Electron Microscopy Sciences, 15710-S) for 15 min at RT and stored at 4 °C until use in sodium azide 0.05% solution (Merck, RTC000068), to prevent bacterial growth.

### 2.4 | Indentation setup and protocol

Mechanical properties of cultured primary astrocytes were measured with a nanoindenter (Pavone, Optics11 Life, Amsterdam, the Netherlands), using an established protocol (Antonovaite et al., 2020). The 24-well plate containing primary astrocytes was placed into the nanoindenter, where it was held at 37 °C. The nanoindenter uses a cantilever-based force sensor equipped with a glass sphere that makes contact with the sample (Antonovaite et al., 2023). The sensor is moved down into the well using a piezo, while the lateral position of the sphere is monitored via an integrated inverted microscope. Upon touching the sample with the sphere, bending of the cantilever is measured via optical fiber and thus applied force and indentation depth are measured in order to extract the Young’s modulus. Single cell indentation measurements were performed once per cell, at a single position next to the nucleus, using a cantilever stiffness of k=0.025 N/m and 3 µm radius spherical tip (see schematic illustration in Figure 1A). All data from a plate was obtained within 3 h after the plate was taken from the incubator. Data was analyzed in Matlab by fitting initial load-indentation curve (<1 µm depth) with Hertz model to extract Young’s modulus E (Antonovaite et al., 2020).

**Figure 1.**
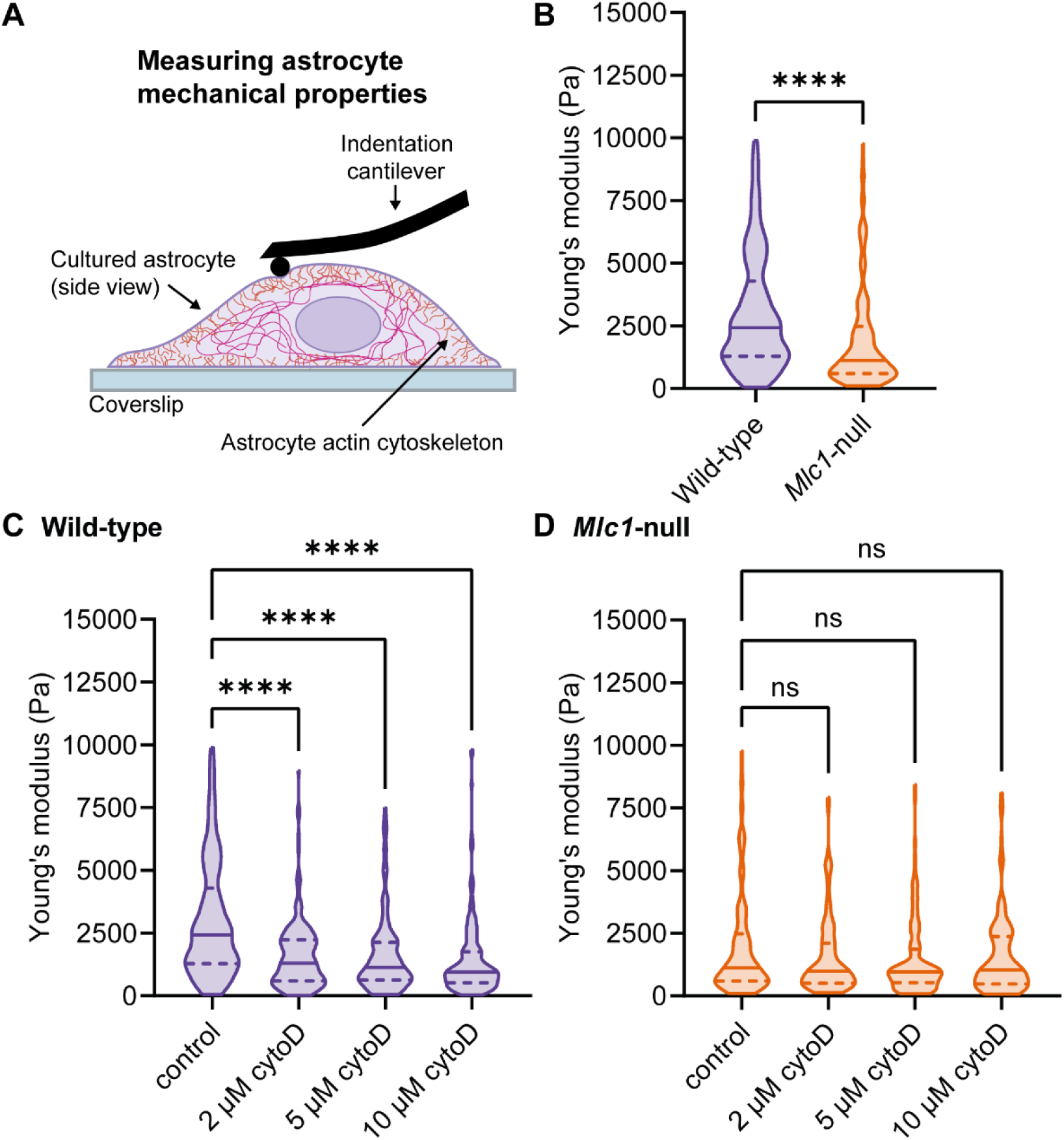
Characterization of mechanical properties of primary wild-type and Mlc1-null astrocytes reveal that Mlc1-null astrocytes are softer than wild-type astrocytes. **A.** Schematic representation of the experimental indentation setup. B. Single cell indentation measurements of wild-type astrocytes (purple) and Mlc1-null astrocytes (orange) in Young’s modulus maps, in pascal (Pa) (wild-type: 2422 (2998) Pa, n=273/N=3, Mlc1-null: 1121 (1181.2) Pa, n=184/N=3, adj. P<0.0001). C,D. Modulating the cytoskeleton with an actin polymerization inhibitor, cytochalasin D (cytoD; 2 µM, 5 µM or 10 µM) in wild-type (C; wild-type control: 2422 (2998) Pa, n=273/N=3, wild-type 2 µM: 1293 (1643.2) Pa, n=178/N=3, adj. P<0.0001, wild-type 5 µM: 1135 (1497) Pa, n=175/N=3, adj. P<0.0001, wild-type 10 µM: 942 (1241.5) Pa, n=200/N=3, adj. P<0.0001), and in Mlc1-null astrocytes (D; Mlc1-null control: 1121 (1881.2) Pa, n=184/N=3, Mlc1-null 2 µM: 987 (1603) Pa, n=127/N=3, adj. P=0.6872, Mlc1-null 5 µM: 963.5 (1329.5) Pa, n=128/N=3, adj. P=0.1727, Mlc1-null 10 µM: 1037 (1896) Pa, n=129/N=3, adj. P=0.5601). N = 3 mice per genotype. All data are presented as median and interquartile range (IQR) values.

### 2.5 | Mass spectrometry

Total protein extracts derived from wild-type and *Mlc1*-null astrocyte pellets (prepared as described in section 2.3) were performed as previously described (Lanciotti et al., 2012). Briefly, pellets were lysed in a buffer containing 1% Triton X-100, 0.5% sodium deoxycholate, 150 mM NaCl, 10 mM HEPES (pH 7.4) and protease (Sigma-Aldrich, P8340) and phosphatase (Roche Diagnostics, Mannheim, Germany) inhibitor cocktail. Lysates were passed through a 26-gauge needle 15 times, incubated on ice for 20 min and centrifuged at 14.000 rpm for 20 min at 4 °C. Protein loading content was quantified using a bicinchoninic acid (BCA) protein assay kit (Thermo Fisher Scientific, Italy). Samples obtained were analyzed in triplicate: 50 µg of cell lysates derived from each sample were sonicated in an ultrasonic bath for 20 min, incubated in 30 mM Tris(2-carboxylethyl)phosphine (TCEP) for 10 min and then in 20 mM iodoacetamide for 30 min in the dark at RT. 200 µl of acetone, methanol, and ethanol mixture (25/25/50) was added and the proteins were precipitated ON at −20 °C. The protein pellet was centrifuged (14000 x *g*, 15 min) and digested with 12 ng/µl trypsin (Promega Corporation, WI, United States) in 25 mM ammonium bicarbonate and 1 M urea at 37 °C ON. Formic acid was added to a final concentration of 0.1% to stop the enzymatic digestion and an aliquot of the resulting peptide solution was injected in an Ultimate 3000 UHPLC (Dionex, Thermo Fisher Scientific Waltham, Massachusetts, USA) coupled with an Orbitrap Fusion Tribrid mass spectrometer (Thermo Fisher Scientific, Waltham, Massachusetts, USA). Peptides were desalted on a trap column (Acclaim PepMap 100 C18, Thermo Fisher, Scientific Waltham, Massachusetts, USA) and then separated on a 45-cm-long silica capillary (Silica Tips FS 360-75-8, New Objective, MA, United States), packed in-house with a C18, 1.9 μm, 100 Å resin (Michrom BioResources, CA, United States).

The analytical separation ran for 180 min using a gradient of buffer A (5% acetonitrile and 0.1% formic acid) and buffer B (95% acetonitrile and 0.1% formic acid). The gradient started with 5% of buffer B, which rose to 6% in 5 min, to 32% in 130 min, to 55% in 25 min, and up to 80% in 4 min and then washing and equilibrating steps were added until a total 180 min run time. Full-scan MS data were acquired in the 350–1,550 m/z mass range in the Orbitrap with 120K resolution. Data-dependent acquisition was performed in top-speed mode (3 s long maximum total cycle): the most intense precursors with charge >1 were selected through a monoisotopic precursor selection (MIPS) filter, quadrupole isolated and fragmented by 30% higher-energy collision dissociation (HCD). Fragment ions were analyzed in the ion trap with rapid scan rate. Raw data were analyzed by the software Proteome Discoverer 2.4 (Thermo Fisher Scientific) using the reviewed version of UniProtKB/Swiss-Prot mouse (taxonomy 10090) database containing 17090 sequences. Spectral matches were filtered using Percolator node, with 1% false discovery rate (FDR) based on q values. Only master proteins were considered and specific trypsin cleavages with two miss-cleavages were admitted. Cysteine carbamydomethylation was set as static modification, while methionine oxidation and N-acetylation on protein terminus were set as variable modifications. 10 ppm and 0.6 Da tolerance were considered for MS and MS/MS data assignment, respectively. Quantification was based on precursor ion intensity of unique and razor peptides and abundance values were normalized on total peptide amount.

Quantitative results were analyzed by Proteome Discoverer: protein ratio calculation was based on pairwise ratio. Proteins were filtered considering only those quantified at least twice in one group. Of this group, proteins were considered up- or downregulated in *Mlc1*-null *vs* wild-type if log2 ratio *Mlc1*-null/wild-type was greater than 0.6 or lower than −0.6, respectively, with an adjusted P-value of < 0.05. Principal component analysis (PCA) was obtained using SRplot (Tang et al., 2023), including only proteins detected in all samples (n=5205). Biological process of enrichment analysis on differentially-modulated proteins was conducted by using the WEB-based GEne SeT AnaLysis Toolkit (Webgestalt)(Liao et al., 2019), looking at the top 10 level categories enriched with 0.05 FDR evaluated by Benjamini correction. Proteins involved in cytoskeleton organization based on Webgestalt analysis were analyzed by STRING (Szklarczyk et al., 2023), and clustered with the Markov Cluster Algorithm (Van Dongen, 2008). All the mass spectrometry proteomics data have been deposited to the ProteomeXchange Consortium via the MassIVE (Mass Spectrometry Interactive Virtual Environment) with the identifier MSV000097012 (Computer Science and Engineering, 2025).

### 2.6 | Immunocytochemistry and imaging

Plates with fixed primary astrocytes on coverslips (see 2.3 for preparation) were taken to RT. After removing the 0.05% sodium azide in the fume hood, coverslips were washed 3 times with DPBS, taken out of the fume hood and washed again 3 times with DPBS. After incubation in blocking buffer (DPBS + 5% Normal Goat Serum (NGS; Gibco^TM^, 16210-064) + 0.1% Bovine Serum Albumin (BSA; Sigma-Aldrich, A4919) + 0.3% Triton X-100 (Sigma-Aldrich, X100)) for 1 h at RT for permeabilization and to block nonspecific binding, coverslips were incubated with primary vinculin antibody and fluorescently-labelled F-actin probe diluted in blocking buffer (4 °C, ON). The next day coverslips were washed 3 times with DPBS and stained with secondary antibody diluted in blocking buffer for 1 h at RT. Coverslips were then washed 3 times with DPBS, and with the third wash stained for 4’,6-diamidino-2-phenylindole (DAPI; 1:2000; Sigma, D9542, 5 mg/ml) diluted in DPBS. Coverslips were washed once with DPBS to remove excess DAPI and mounted on glass slides using ProLong^TM^ Glass Antifade Mountant mounting medium (Invitrogen, P36984), then, in the dark, stored first at RT for 24 h and then at 4 °C until use. The following antibodies were used: anti-vinculin (1:800; Sigma-Aldrich, V9131, mouse monoclonal antibody IgG1, 5-10 mg/mL), and, as secondary, Alexa Fluor^TM^ 488 goat anti-mouse IgG (1:1000; Invitrogen, A-11001, 2 mg/mL). For staining of F-actin, a tetramethylrhodamine isothiocyanate (TRITC)-conjugated phalloidin probe (1 unit; Invitrogen, R415, 300 units) was used. Multiple Z-stacks with a z step size of 0.225 µm were obtained on an inverted Nikon Eclipse Ti2 confocal microscope (Nikon, Japan) with an oil-immersion objective (Nikon Plan Fluor 40X/NA 1.3/ WD 240 µm) and intermediate magnification of 1.5X. DAPI was excited at 406 nm (emission filter: 425-475), vinculin at 488 nm (emission filter: 500-550) and F-actin at 561 nm (emission filter: 570-616).

### 2.7 | Imaging analysis

Images were analyzed using Fiji (Schindelin et al., 2012). Summation (SUM) and maximum (MAX) intensity projections were made, and cell regions of interest (ROIs) were drawn based on the F-actin channel with the wand tracing tool. Additional square background ROIs were drawn for each image on a region where no specific staining was observed for any of the channels. ROIs were measured for all channels with the multi measure tool to obtain morphological and other parameters mentioned below. Values were background-corrected by subtracting the fluorescence intensity from a background square for each corresponding image. Cells with an area larger than 10000 µm^2^ were excluded.

Vinculin was used as a general marker for FAs. FA analysis was done following a modified protocol (Güler et al., 2021). MAX intensity projections were split into different channels, and the channel containing the vinculin staining was adjusted into 8-bit. The image was processed by subtracting the background with Fast Fourier Transformation (FFT) Bandpass Filter. The images were thresholded with the Huang method. FAs were counted and analyzed with the ‘Analyze Particles’ macro (size 10-Infinity in pixels, circularity 0.00-0.99), with the cell ROI selected to only include FAs in the cell of interest. Thresholding alters fluorescence intensity levels of the image; therefore, the generated ROIs of FAs were applied to the original 16-bit SUM intensity projection image to determine fluorescence intensity values of all individual FAs. These values were also background-corrected as described previously, and values were averaged per cell.

All fluorescence intensity values were obtained from SUM intensity projections and normalized to averaged values of wild-type cells imaged on the same days. All data was obtained from cells from 3 different mice per genotype. Based on differences in area, data was grouped into three separate groups: 0 – 1500 µm^2^, 1500 – 4000 µm^2^ and 4000+ µm^2^. Each group contains cells from three different mice.

### 2.8 | Statistics

Data representation and statistical analysis were performed using GraphPad Prism version 10.2.0 for Windows (GraphPad Software, Boston, Massachusetts USA, www.graphpad.com). Data were tested for normality with a Kolmogorov-Smirnov test. Statistical analysis of indentation data was done with a Mann-Whitney *U* test (Figure 1B), and a Kruskal-Wallis test with Dunn’s multiple comparisons test, by comparing all groups with the control group (Figure 1C and 1D). Statistical analysis of the imaging data was done with a Mann-Whitney *U* test with a Holm-Šidák multiple comparisons test, by comparing wild-type to *Mlc1*-null astrocytes in every size group (0 – 1500 µm^2^, 1500 – 4000 µm^2^ and 4000+ µm^2^; Figure 3 and 5). Cumulative distributions of the area of wild-type and *Mlc1*-null astrocytes were compared with a Kolmogorov-Smirnov test (Figure 3B). Indentation data was reported as median (interquartile range [IQR]); all other data are expressed as mean ± SEM. Statistically significant differences were defined as *P*≤0.05.

## 3 | RESULTS

### 3.1 | Mlc1-null astrocytes have altered mechanical properties

We investigated mechanical properties of cultured primary *Mlc1*-null and wild-type astrocytes with indentation measurements. To minimize the influence of the plastic substrate on indentation measurements, the chosen measurement point for each cell was close to the nucleus, at its thickest point (Figure 1A) (Antonovaite et al., 2020). To determine cell stiffness, the Young’s modulus, or elastic modulus, was determined for individual cells. This measure reflects the resistance of a substance to elastic deformation. *Mlc1* -null astrocytes had a significantly lower Young’s modulus than wild-type astrocytes (Figure 1B), indicating that loss of MLC1 leads to softer astrocytes. To investigate how disruption of the astrocyte actin cytoskeleton would influence cell stiffness, wild-type and *Mlc1*-null astrocytes were treated with an increasing dose of the actin polymerization inhibitor cytochalasin D. Disrupting the actin cytoskeleton caused a significant decrease in the Young’s modulus of wild-type astrocytes for all three used concentrations of cytochalasin D (Figure 1C). Thus, actin cytoskeleton disruption led to softening of wild-type astrocytes. Treatment of *Mlc1*-null astrocytes with cytochalasin D did not induce a further decrease in the Young’s modulus (Figure 1D). This suggests that the already impaired structural integrity of *Mlc1*-null astrocytes could not be exacerbated by disruption of the actin cytoskeleton. In conclusion, indentation measurements revealed that *Mlc1*-null astrocytes are softer than wild-type astrocytes, and that disruption of the actin cytoskeleton in *Mlc1*-null astrocytes does not lead to additional softening.

### 3.2 | Proteomic analysis reveals alterations of cytoskeleton-related pathways in Mlc1-null astrocytes

We characterized the proteome of cultured primary wild-type and *Mlc1*-null astrocytes. Principal Component Analysis (PCA) showed a clear separation between wild-type and *Mlc1*-null samples (Figure 2A). MLC1 was only detected in wild-type samples, as expected (Figure S1). Proteomic analysis led to the identification of 6232 proteins, of which 96% (5960 proteins) could be quantified (for the total list see Table S1). After data filtering, 5858 proteins were analyzed: 125 proteins were identified as downregulated and 134 as upregulated in *Mlc1*-null samples compared to wild-type samples (Table S2; Figure 2B). These up- and downregulated proteins are shown in a volcano plot with several cytoskeleton-related proteins and other proteins of interest indicated (Figure 2C). Analysis of biological pathways showed significant enrichment for pathways related to the cytoskeleton: microtubule bundle formation, microtubule-based process and cytoskeleton organization (Figure 2D; in bold). Proteins from the biological process cytoskeleton organization were further analyzed for protein-protein interaction networks (Figure 2E). As part of the cytoskeleton organization cluster, glial fibrillary acidic protein (GFAP) was downregulated, and the microtubule associated protein 1B (MAP1B) was upregulated in *Mlc1*-null astrocytes. The actin protein ACTG1 was downregulated and myosin protein MYO1C was upregulated. In conclusion, several cytoskeleton-related pathways are found to be differentially regulated in *Mlc1*-null astrocytes.

**Figure 2.**
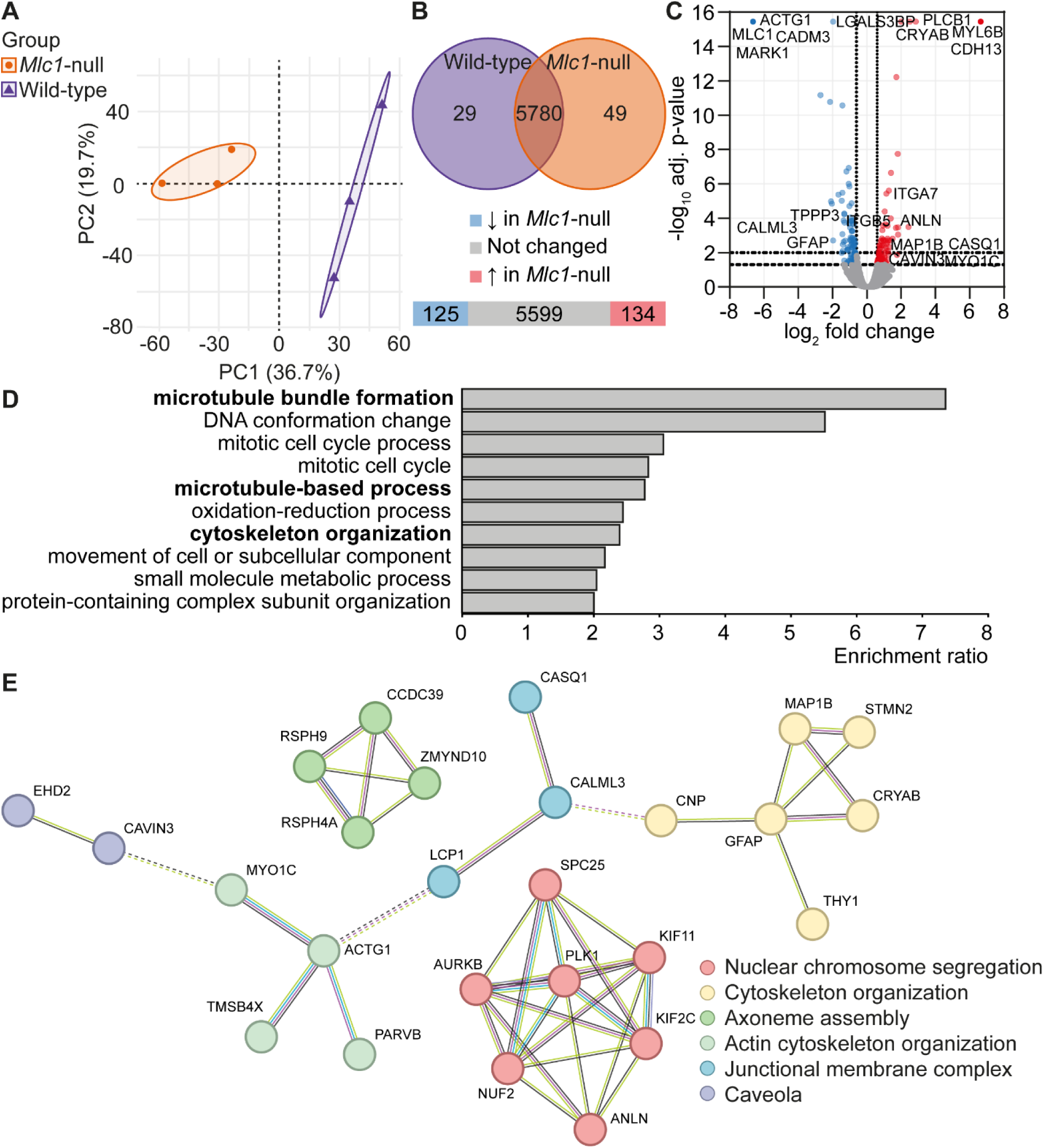
Proteomic analysis shows enrichment of cytoskeleton-related pathways in Mlc1-null astrocytes. **A.** Principal component analysis (PCA) on proteins quantified in all samples (n=5205) shows a clear separation between primary wild-type (purple) and Mlc1-null (orange) astrocytes. The first component (PC1) separates wild-type from Mlc1-null astrocytes and explains 36.7% of the variability. The second component (PC2) is representing 19.7% of the variability. **B,C.** Venn diagram shows total number of 5858 proteins quantified at least twice in one group. Of these 5858 proteins, 5780 were detected in both wild-type (purple) and Mlc1-null (orange) astrocytes, 29 only in wild-type and 49 only in Mlc1-null astrocytes. Differentially expressed proteins in Mlc1-null astrocytes compared to wild-type astrocytes, based on a fold change > 0.06 or < −0.06 with an adj. P<0.05, are shown in the bar plot **(B)** and the volcano plot **(C)**. Bar plot shows the number of proteins not changed (n=5599; grey), down-(n=125; blue) and upregulated (n=134; red). In the volcano plot, MLC1 and several other proteins related to the cytoskeleton are highlighted. **D.** The WEB-based GEne SeT AnaLysis Toolkit was used for biological process of enrichment analysis on differentially modulated proteins. The top 10 categories that are enriched with a False Discovery Rate (FDR) of ≤0.05 are shown. Three biological processes related to the cytoskeleton are highlighted in bold: microtubule bundle formation, microtubule-based process, and cytoskeleton organization. E. Analysis of protein-protein interaction networks on proteins involved in cytoskeleton organization. N = 3 mice per genotype.

### 3.3 | Mlc1-null astrocytes show no differences in morphology or F-actin fluorescence intensity

To investigate the organization of the actin cytoskeleton in cultured primary wild-type and *Mlc1*-null astrocytes, we performed immunofluorescence staining of F-actin. This allowed us to also visualize general cell morphology (Figure 3A). Primary astrocytes were heterogeneous in shape and morphology. On average, wild-type and *Mlc1*-null astrocytes span the same area. A cumulative distribution analysis of cell area of wild-type and *Mlc1*-null astrocytes showed that about half of the cells were sized between 1500 and 4000 µm^2^ (Figure 3B). We did observe a non-significant trend towards an overrepresentation of smaller cells in astrocytes isolated from *Mlc1*-null mice. Since the organization of the cytoskeleton strongly depends on total cell size, we split the cells into three distinct size categories for further analysis. The perimeter and the aspect ratio (ratio of width and height) were similar between wild-type and *Mlc1*-null astrocytes in all three categories, indicating comparable morphology (Figure 3C and 3D). Fluorescence intensity of F-actin showed no significant differences between wild-type and *Mlc1*-null astrocytes. The group of small (0 – 1500 µm^2^) *Mlc*1-null astrocytes showed a trend towards lower F-actin fluorescence levels (Figure 3E). We did not observe any obvious alterations in the distribution of F-actin throughout the astrocytes. Overall, immunofluorescence analysis suggested no strong alterations in the levels or organization of F-actin between wild-type and *Mlc1*-null astrocytes.

**Figure 3.**
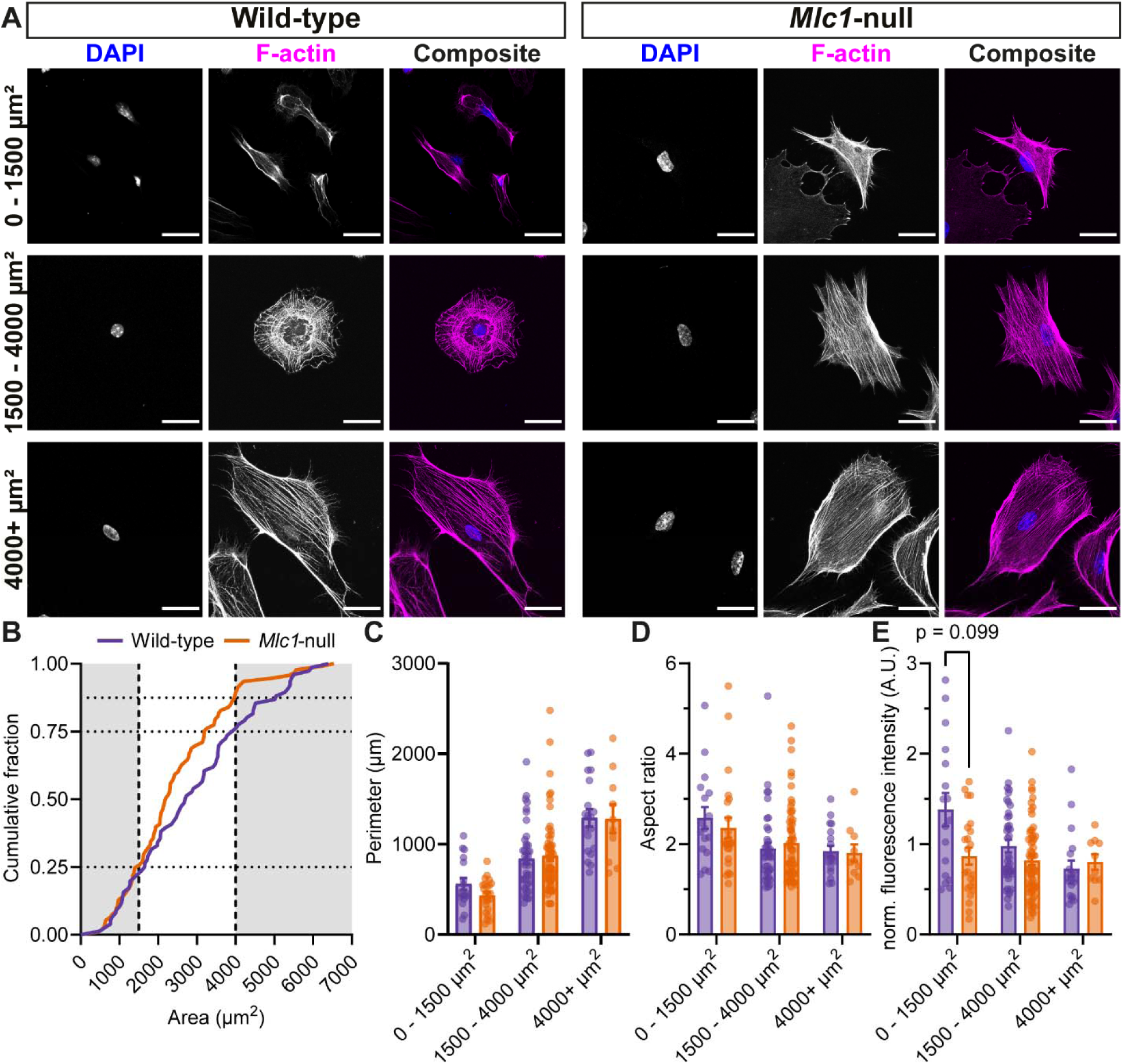
Mlc1-null astrocytes have similar morphological parameters to wild-type astrocytes, and cells of size 1500 – 4000 µm^2^ show a trend in decrease of F-actin fluorescence intensity. **A.** Representative confocal images of primary wild-type and Mlc1-null astrocytes, stained for the nucleus with DAPI (blue) and F-actin (magenta). The cells are split in three different size groups: 0 – 1500 µm^2^, 1500 – 4000 µm^2^ and 4000+ µm^2^. Scale bars: 30 µm. **B.** Cumulative distribution function of the area of wild-type (purple) and Mlc1-null (orange) astrocytes (D=0.1773, approx. P=0.1443). **C,D**. Cell morphology parameters perimeter (**C**) and aspect ratio (**D**; ratio between height and width of the cells) of wild-type (purple) and Mlc1-null (orange) astrocytes (perimeter: wild-type 0 – 1500 µm^2^: 560.6±65.6 µm, Mlc1-null 0 – 1500 µm^2^: 432.5±37.5 µm, adj. P=0.603, wild-type 1500 – 4000 µm^2^: 841.1±58.6 µm, Mlc1-null 1500 – 4000 µm^2^: 873.2±53.3 µm, adj. P=0.902, wild-type 4000+ µm^2^: 1292.7±96.1 µm, Mlc1-null 4000+ µm^2^: 1281.8±157.1 µm, adj. P=0.902; aspect ratio: wild-type 0 – 1500 µm^2^: 2.58±0.24, Mlc1-null 0 – 1500 µm^2^: 2.36±0.23, adj. P=0.584, wild-type 1500 – 4000 µm^2^: 1.90±0.13, Mlc1-null 1500 – 4000 µm^2^: 2.03±0.11, adj. P=0.753, wild-type 4000+ µm^2^: 1.85±0.12, Mlc1-null 4000+ µm^2^: 1.80±0.19, adj. P=0.753). E. Normalized and background-corrected fluorescence intensity of F-actin in wild-type (purple) and Mlc1-null (orange) astrocytes (wild-type 0 – 1500 µm^2^: 1.38±0.18, Mlc1-null 0 – 1500 µm^2^: 0.87±0.09, adj. P=0.099, wild-type 1500 – 4000 µm^2^: 0.98±0.07, Mlc1-null 1500 – 4000 µm^2^: 0.82±0.05, adj. P=0.15, wild-type 4000+ µm^2^: 0.73±0.09, Mlc1-null 4000+ µm^2^: 0.80±0.08, adj. P=0.266). Each group contains cells from three different mice per genotype (wild-type 0 – 1500 µm^2^: n=17/N=3, Mlc1-null 0 – 1500 µm^2^: n=23/N=3, wild-type 1500 – 4000 µm^2^: n=40/N=3, Mlc1-null 1500 – 4000 µm^2^: n=60/N=3, wild-type 4000+ µm^2^: n=19/N=3, Mlc1-null 4000+ µm^2^: n=10/N=3). All data is presented as mean ± SEM, with dots indicating individual values of cells in bar graphs.

### 3.4 | Alterations in focal adhesions in Mlc1-null astrocytes

Patch-clamp studies on *Mlc1*-null astrocytes suggested that the cell membrane of these cells was loosely connected to the cytoskeleton. The membrane was pulled further into the patch-clamp pipette when trying to apply suction, compared to wild-type astrocytes (Bisseling and Min, personal communication). Therefore, we studied whether the connection of cytoskeleton to membrane was altered in *Mlc1*-null astrocytes. FA complexes connect the actin cytoskeleton to the cell membrane and the extracellular matrix (ECM) through integrin receptors, and can be visualized by immunofluorescence staining for the FA marker vinculin (Burridge et al., 1988). Co-staining of F-actin (magenta) and vinculin (green) revealed their partial colocalization in white (Figure 4). In the intermediately-sized cell group (1500 – 4000 µm^2^), fluorescence intensity of vinculin was lower in *Mlc1*-null astrocytes when compared to wild-type astrocytes (Figure 5A). We quantified the number, morphology and fluorescence intensity of individual FAs by using a particle analysis algorithm (Figure 5C-F; see Figure 5B for examples analysis) (Güler et al., 2021). Compared to wild-type astrocytes, the number of FAs per µm^2^ was 30% lower in the largest *Mlc1*-null astrocytes (4000+ µm^2^; Figure 5C). In this size category we did not observe a change in the percentage of the cell covered by FAs (Figure 5D), which can be explained by a combination of a decrease in number of FAs, and an increase in FA size (Figure 5E). There was no difference in the fluorescence intensity of FAs between wild-type and *Mlc1*-null astrocytes (Figure 5F). Together, our findings illustrate several alterations in the properties and morphology of FAs in primary astrocytes from *Mlc1*-null mice.

**Figure 4.**
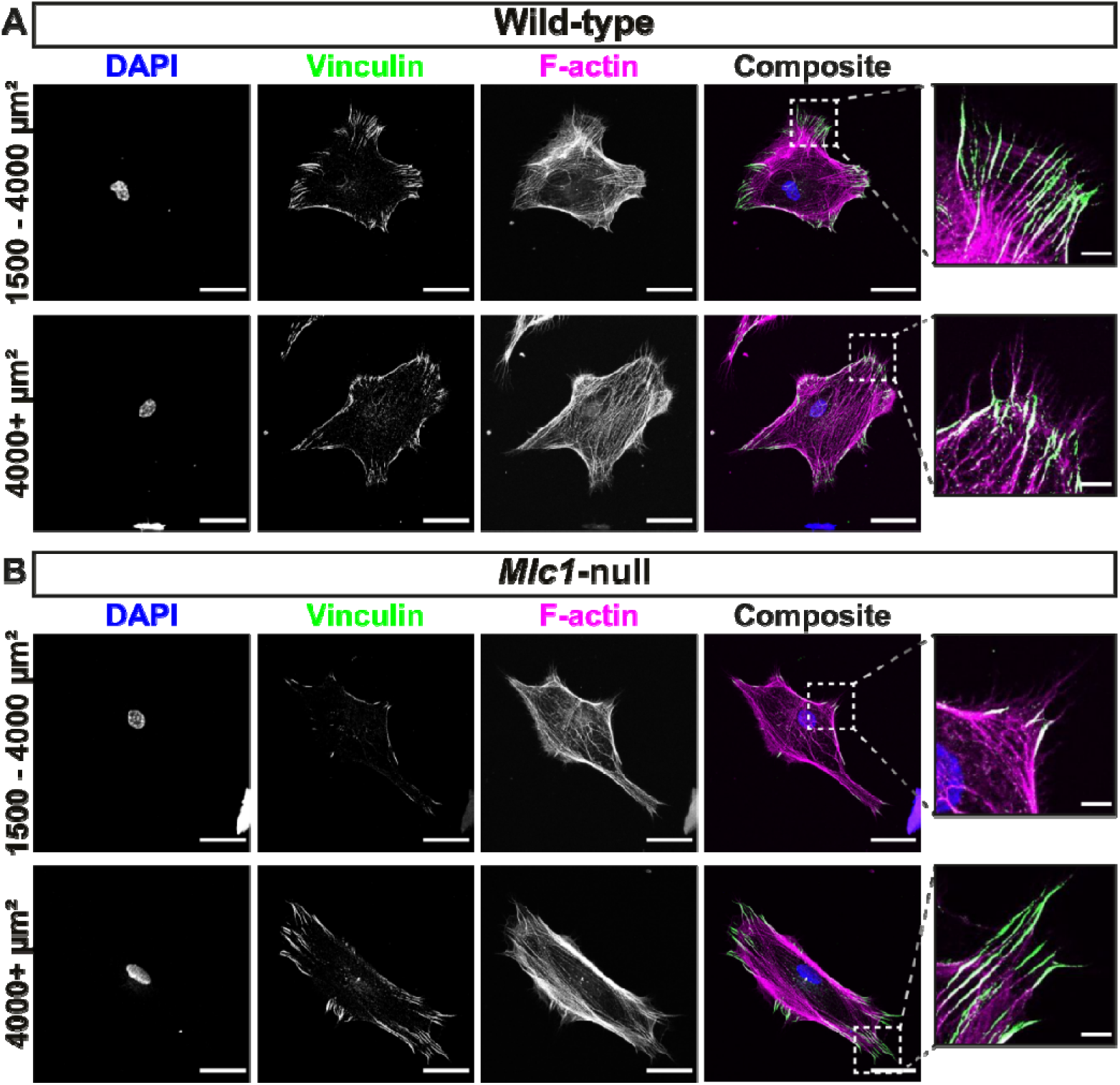
Immunofluorescence staining of vinculin, a focal adhesion protein, in wild-type and Mlc1-null astrocytes. Representative confocal images of primary wild-type **(A)** and Mlc1-null **(B)** astrocytes in size groups 1500 – 4000 µm^2^ and 4000+ µm^2^, stained for DAPI (blue), vinculin (green) and F-actin (magenta). Focal adhesions are shown in the enlarged images. Scale bars: 30 µm, and 5 µm for the enlarged images.

**Figure 5.**
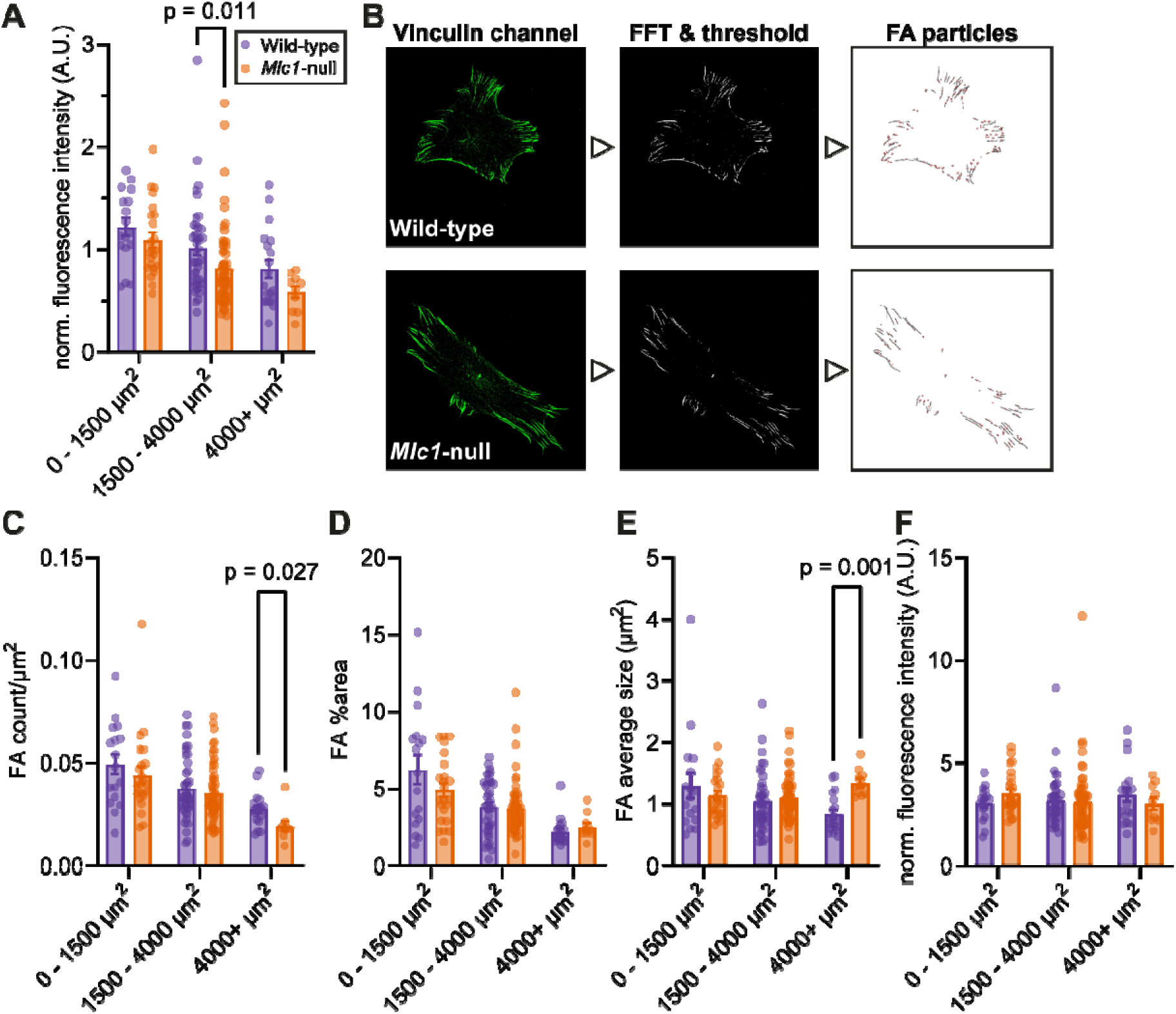
Focal adhesions, represented by vinculin, are altered in Mlc1-null astrocytes. **A.** Normalized and background-corrected fluorescence intensity of vinculin in wild-type (purple) and Mlc1-null (orange) astrocytes in size groups 0 – 1500 µm^2^, 1500 – 4000 µm^2^ and 4000+ µm^2^ (wild-type 0 – 1500 µm^2^: 1.22±0.09, Mlc1-null 0 – 1500 µm^2^: 1.09±0.08, adj. P=0.317, wild-type 1500 – 4000 µm^2^: 1.01±0.07, Mlc1-null 1500 – 4000 µm^2^: 0.81±0.05, adj. P=0.011, wild-type 4000+ µm^2^: 0.81±0.09, Mlc1-null 4000+ µm^2^: 0.58±0.06, adj. P=0.317). **B.** Examples of particle analysis in a wild-type (first row) and Mlc1-null (second row) astrocyte. Starting with the original MAX projection of the vinculin channel, then the same image after background correction and thresholding. The obtained ROIs of all the FAs in the cell (third image) were used to extract different FA parameters. **C-F.** Parameters of FA in wild-type (purple) and Mlc1-null (orange) astrocytes in size groups 0 – 1500 µm^2^, 1500 – 4000 µm^2^ and 4000+ µm^2^. **C.** The number of FAs in the cell per µm^2^ (wild-type 0 – 1500 µm^2^: 0.049±0.005, Mlc1-null 0 – 1500 µm^2^: 0.044±0.004, adj. P=0.585, wild-type 1500 – 4000 µm^2^: 0.038±0.002, Mlc1-null 1500 – 4000 µm^2^: 0.035±0.002, adj. P=0.585, wild-type 4000+ µm^2^: 0.027±0.002, Mlc1-null 4000+ µm^2^: 0.019±0.002, adj. P=0.027). **D.** The percentage of the cell area that contains FAs (wild-type 0 – 1500 µm^2^: 6.23±0.95 %, Mlc1-null 0 – 1500 µm^2^: 4.90±0.45 %, adj. P=0.695, wild-type 1500 – 4000 µm^2^: 3.78±0.28 %, Mlc1-null 1500 – 4000 µm^2^: 3.71±0.22 %, adj. P=0.695, wild-type 4000+ µm^2^: 2.17±0.2 %, Mlc1-null 4000+ µm^2^: 2.51±0.26 %, adj. P=0.417). **E.** The average size of FAs per cell (wild-type 0 – 1500 µm^2^: 1.29±0.21 µm^2^, Mlc1-null 0 – 1500 µm^2^: 1.14±0.07 µm^2^, adj. P=0.914, wild-type 1500 – 4000 µm^2^: 1.05±0.08 µm^2^, Mlc1-null 1500 – 4000 µm^2^: 1.10±0.04 µm^2^, adj. P=0.479, wild-type 4000+ µm^2^: 0.83±0.07 µm^2^, Mlc1-null 4000+ µm^2^: 1.34±0.07 µm^2^, adj. P=0.001). **F.** Normalized and background-corrected fluorescence intensity of FAs. Individual FA values were averaged per cell (wild-type 0 – 1500 µm^2^: 3.04±0.2, Mlc1-null 0 – 1500 µm^2^: 3.5±0.22, adj. P=0.832, wild-type 1500 – 4000 µm^2^: 3.17±0.2, Mlc1-null 1500 – 4000 µm^2^: 3.1±0.21, adj. P=0.832, wild-type 4000+ µm^2^: 3.45±0.32, Mlc1-null 4000+ µm^2^: 3±0.3, adj. P=0.832). Each group contains cells from three different mice per genotype (wild-type 0 – 1500 µm^2^: n=17/N=3, Mlc1-null 0 – 1500 µm^2^: n=23/N=3, wild-type 1500 – 4000 µm^2^: n=40/N=3, Mlc1-null 1500 – 4000 µm^2^: n=60/N=3, wild-type 4000+ µm^2^: n=19/N=3, Mlc1-null 4000+ µm^2^: n=10/N=3). All data is presented as mean ± SEM, with dots indicating individual values of cells in bar graphs.

## 4 | DISCUSSION

Dysfunctional astrocytes cause disturbed ion and water homeostasis in MLC, but the exact mechanism of this disturbance is not fully understood. Here, we use different approaches to demonstrate that the mechanical properties of primary astrocytes are altered upon loss of MLC1. Using a specialized indentation method, we show that *Mlc1*-null astrocytes are softer than wild-type astrocytes. Proteomic analysis of *Mlc1*-null astrocytes reveals alterations in several cytoskeleton-related pathways. Confocal imaging of cytoskeletal organization in *Mlc1-* null astrocytes indicates no changes in actin cytoskeleton structure. Instead, we observe alterations in FAs in *Mlc1*-null astrocytes, suggesting that the interaction between cytoskeleton, cell membrane and ECM is altered.

We use indentation experiments to show that *Mlc1*-null astrocytes are soft. Cytochalasin D (Brenner & Korn, 1979), an actin polymerization inhibitor that causes a near-complete collapse of the actin cytoskeleton, makes wild-type astrocytes softer but does not induce further softening in *Mlc1*-null astrocytes. Since actin is the main contributor to cell stiffness in astrocytes (Curry et al., 2017), we investigate whether reduced cell stiffness in *Mlc1*-null astrocytes could be due to disruption of the actin cytoskeleton. However, imaging shows that the actin cytoskeleton is morphologically intact in *Mlc1*-null astrocytes.

Proteomic analysis reveals that *Mlc1* -null astrocytes show differential expression of proteins implicated in cytoskeleton-related pathways. In addition to proteins related to the actin cytoskeleton, several microtubule-related proteins are altered. Microtubules are involved in intracellular transport, cell division and structural support (Weigel et al., 2021), and play a role in mechanotransduction (Seetharaman et al., 2022). Since disturbed mechanotransduction could be implicated in MLC, future studies on microtubule alterations in MLC are warranted.

The question remains what causes the softening of *Mlc1*-null astrocytes. Whilst performing patch-clamp experiments, we noticed that the membranes of *Mlc1*-null astrocytes were pulled further into the pipette when generating a gigaseal (Bisseling and Min, personal communication). These observations could not be quantified but suggest a compromised attachment of the membrane to the cytoskeleton. This prompted us to investigate cytoskeleton-membrane-ECM interactions. We focused on FAs, large protein complexes that connect the actin cytoskeleton, membrane and ECM (Legerstee & Houtsmuller, 2021). We observed alterations in expression of the FA protein vinculin. Previous studies have established that loss of vinculin reduces cell stiffness and decreases adhesion to the ECM (Goldmann et al., 1998; Mierke et al., 2008; Klemm et al., 2009; Mierke et al., 2010). Changes in FAs in *Mlc1*-null astrocytes might therefore lead to a reduced tensile strength on the actin cytoskeleton, resulting in softer *Mlc1*-null astrocytes.

FA are highly dynamic structures, and their turnover is essential for cell motility and migration (Yamaguchi & Knaut, 2022). Tension-induced activation of vinculin stabilizes and matures FAs, which enforces cytoskeleton-membrane-ECM interaction and enables cells to endure mechanical tension (Galbraith et al., 2002; Grashoff et al., 2010; Carisey et al., 2013; Dumbauld et al., 2013). As FA dynamics determine the strength of this interaction, an exciting next step would be to compare FA dynamics in wild-type and *Mlc1*-null astrocytes using live-cell imaging. Interestingly, THY-1 was upregulated in *Mlc1*-null astrocytes. This is a glycoprotein that binds integrin receptors in astrocytes and promotes FA formation and cell spreading (Leyton et al., 2001). An explanation might be that FAs are formed but not maintained in *Mlc1*-null astrocytes, leading to high turnover and less stable FAs and an increase in THY-1 levels. Such dynamic changes could be determined with live-cell imaging.

Cultured primary astrocytes are a useful tool to understand the cell biology of MLC1. However, they do not replicate in vivo morphology. It is therefore crucial to explore whether cytoskeletal and mechanical properties of intact astrocytes in the brain are altered in MLC. Of particular interest is studying the mechanical properties of perivascular astrocyte endfeet, where MLC1 is highly expressed (Boor et al., 2005). These structures are anchored to the vasculature through interaction of membrane proteins with the ECM. Our data suggests a compromised cytoskeleton-membrane-ECM interaction upon loss of MLC1. In line with this, an earlier study established that absence of MLC1 leads to disturbed adhesion of endfeet to the vasculature (Gilbert et al., 2021). ECM proteins form the perivascular basal lamina to which the endfoot adheres. Notably, a loss of ECM proteins can lead to brain abnormalities that resemble MLC (Min & van der Knaap, 2018). Pathogenic variants in one such protein, laminin α2, leads to Merosin-deficient Congenital Muscular Dystrophy (CMD) (Helbling-Leclerc et al., 1995), and MRI abnormalities of the brain white matter in these patients display striking similarities with MLC (Philpot et al., 1995; van der Knaap et al., 1997). The DAGC, which binds laminin in the ECM, is colocalized with MLC1 in astrocyte endfeet (Boor et al., 2007; Ambrosini et al., 2008). A specialized structure known as the costamere, composed of the DAGC and FAs, is present in striated muscle cells (where MLC1 is not expressed), where it regulates cytoskeleton-membrane-ECM interactions (Jaka et al., 2015). We hypothesize that MLC1 is part of an analogous structure in astrocyte endfeet that regulates these interactions. Future studies to validate such a structure in perivascular endfeet in MLC mouse tissue, with super-resolution imaging, could provide valuable insights into MLC pathology, as well as astrocyte physiology.

An open question is how disrupted mechanical properties of astrocyte endfeet leads to brain edema. Many studies on MLC have provided evidence for (mechanosensitive) ion channel dysfunction and disturbed volume regulation in MLC (Ridder et al., 2011; Jeworutzki et al., 2012; Lanciotti et al., 2012; Passchier et al., 2023). The function of ion and water channels involved in volume regulation is very much dependent on cytoskeleton-membrane-ECM interactions. For example, VRAC is modulated by membrane stretch and the cytoskeleton (Byfield et al., 2006), and chloride currents similar to VRAC activation can be evoked by stretching of integrin receptors (Browe & Baumgarten, 2006). TRPV4 is another important mechanosensitive ion channel implicated in astrocyte volume regulation (Benfenati et al., 2007), and its function strongly depends on its association with the cytoskeleton (Becker et al., 2009). Finally, AQP4 localization is modulated by the actin cytoskeleton (Nicchia et al., 2008). It is likely that alterations in astrocyte mechanical properties contribute to dysregulation of ion channels and thereby lead to defective volume regulation in MLC. In addition, astrocyte endfeet form a crucial component of the so-called glymphatic system (Jessen et al., 2015). Recent computational studies have shown that endfeet might act as ‘valves’ around the vasculature to regulate perivascular fluid flow (Bork et al., 2023; Gan et al., 2023; Gan et al., 2024). Thus, a disruption of the structural integrity of endfeet in MLC might directly alter glymphatic fluid flow by interfering with such a valve function.

Interplay between MLC1 or GlialCAM and the cytoskeleton is also observed outside of astrocyte endfeet. Astrocyte specializations that point towards the synapse, so-called perisynaptic processes, also contain MLC1. In *Mlc1*-null mice, these processes are retracted from synapses due to a shortened tip length (Kater et al., 2023). This phenotype resembles what is seen in astrocytes lacking ezrin (Badia-Soteras et al., 2023), an actin-membrane linker that regulates FA dynamics (Hoskin et al., 2015). MLC1 and GlialCAM have been shown to regulate cell motility and proliferation in various cell lines, and dysregulation of these proteins, as well as their newly identified interaction partner GPRC5B (Alonso-Gardón et al., 2021; Passchier et al., 2023), has been implicated in cancer (Lanciotti et al., 2016; Hwang et al., 2019; Lattier et al., 2020; Kanamori et al., 2023; Wu et al., 2024). MLC1 can regulate actin dynamics by interacting with the Arp2/3 complex (Hwang et al., 2019), which drives lamellipodia protrusion through actin networks advancing at the membrane (Swaminathan et al., 2016; Gautreau et al., 2022). GlialCAM is essential for cell-ECM adhesion (Moh et al., 2005), and regulates FA signaling pathways in glioma (De et al., 2022). These studies show that MLC1 regulates cytoskeleton-related processes in several physiological and pathological conditions.

In conclusion, our study provides a new perspective on astrocyte dysfunction in MLC. We present evidence for a role of MLC1 in regulating astrocyte mechanobiology. Our results provide a basis for future investigations into how MLC1 is involved in tuning of mechanical properties in intact astrocytes, how this relates to astrocyte volume regulation and brain fluid dynamics, and how disruption of these processes leads to chronic brain edema in MLC.

## Supporting information

Supplementary table proteomics

## Acknowledgements

We thank Anastasia Bomhof and Leoni Hoogterp for excellent technical assistance. RM and MSvdK are members of the European Reference Network for Rare Neurological Diseases (ERN-RND), project ID 739510.

