## Supplementary material for "Altered mechanical properties of astrocytes lacking MLC1; implications for the leukodystrophy MLC"

### Supplementary figures

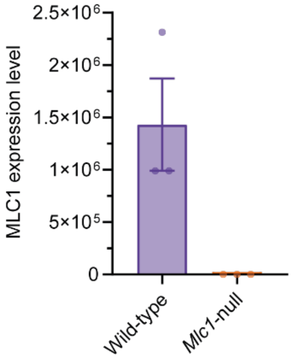

**Figure S1. MLC1 expression in cultured primary wild-type and *Mlc1*-null astrocytes.** Bar plot of MLC1 expression levels in wild-type (purple) and *Mlc1*-null (orange) astrocytes obtained from cell pellets used for proteomic analysis**.** Absence of MLC1 in *Mlc1*-null astrocyte cell pellets was confirmed.

### Supplementary tables

| Up | Accession | Protein description | Down | Accession | Protein description |
|---|---|---|---|---|---|
| ↑ | P14602 | Heat shock protein beta-1 | ↓ | Q07797 | Galectin-3-binding protein |
| ↑ | P23927 | Alpha-crystallin B chain | ↓ | Q9QZQ8 | Core histone macro-H2A.1 |
| ↑ | Q5EBG6 | Heat shock protein beta-6 | ↓ | P20065 | Thymosin beta-4 |
| ↑ | Q9D1C1 | Ubiquitin-conjugating enzyme E2 C | ↓ | P20060 | Beta-hexosaminidase subunit beta |
| ↑ | Q8R2Y2 | Cell surface glycoprotein MUC18 | ↓ | P97429 | Annexin A4 |
| ↑ | Q61738 | Integrin alpha-7 | ↓ | P15105 | Glutamine synthetase |
| ↑ | Q810U4 | Neuronal cell adhesion molecule | ↓ | P31786 | Acyl-CoA-binding protein |
| ↑ | Q3UQA7 | Selenoprotein H | ↓ | Q60963 | Platelet-activating factor acetylhydrolase |

| | | | | | |
|---|---|---|---|---|---|
| ↑ | Q8K298 | Anillin | ↓ | O35683 | NADH dehydrogenase [ubiquinone] 1 alpha subcomplex subunit 1 |
| ↑ | P56390 | Cyclin-dependent kinases regulatory subunit 2 | ↓ | Q923L3 | CUB and sushi domain-containing protein 1 |
| ↑ | P97298 | Pigment epithelium-derived factor | ↓ | P97384 | Annexin A11 |
| ↑ | Q03145 | Ephrin type-A receptor 2 | ↓ | Q8K1A5 | Transmembrane protein 41B |
| ↑ | Q9CPS6 | Adenosine 5'-monophosphoramidase HINT3 | ↓ | Q99P91 | Transmembrane glycoprotein NMB |
| ↑ | Q6P9P6 | Kinesin-like protein KIF11 | ↓ | Q9D6P8 | Calmodulin-like protein 3 |
| ↑ | P63054 | Calmodulin regulator protein PCP4 | ↓ | Q4FK66 | Pre-mRNA-splicing factor 38A |
| ↑ | Q8CH77 | Neuron navigator 1 | ↓ | Q99ML0 | Zinc finger MYND domain-containing protein 10 |
| ↑ | Q99104 | Unconventional myosin-Va | ↓ | Q9ERY9 | Ergosterol biosynthetic protein 28 homolog |
| ↑ | Q8BJZ3 | Protein lifeguard 3 | ↓ | Q9D9V4 | Radial spoke head protein 9 homolog |
| ↑ | Q8C2B3 | Histone deacetylase 7 | ↓ | Q61845 | Meiosis-expressed gene 1 protein |
| ↑ | P18155 | Bifunctional methylenetetrahydrofolate dehydrogenase/cyclohydrolase, mitochondrial | ↓ | Q9DAJ5 | Dynein light chain roadblock-type 2 |
| ↑ | Q8R060 | Protein zwilch homolog | ↓ | P56382 | ATP synthase subunit epsilon, mitochondrial |
| ↑ | P07607 | Thymidylate synthase | ↓ | Q8JZV9 | 3-hydroxybutyrate dehydrogenase type 2 |
| ↑ | Q91XC8 | Death-associated protein 1 | ↓ | Q8BGZ7 | Keratin, type II cytoskeletal 75 |

| | | | | | |
|---|---|---|---|---|---|
| ↑ | P70248 | Unconventional myosin-If | ↓ | Q9CRC6 | BLOC-1-related complex subunit 7 |
| ↑ | O70456 | 14-3-3 protein sigma | ↓ | Q9D620 | Rab11 family-interacting protein 1 |
| ↑ | P04184 | Thymidine kinase, cytosolic | ↓ | Q99J94 | Solute carrier organic anion transporter family member 1A6 |
| ↑ | Q9D600 | DNA replication complex GINS protein PSF2 | ↓ | Q7TQ48 | Sarcalumenin |
| ↑ | O09165 | Calsequestrin-1 | ↓ | Q8BYM7 | Radial spoke head protein 4 homolog A |
| ↑ | Q9DAP7 | Histone chaperone ASF1B | ↓ | Q9D4H2 | GRIP and coiled-coil domain-containing protein 1 |
| ↑ | Q61846 | Maternal embryonic leucine zipper kinase | ↓ | Q9D6M3 | Mitochondrial glutamate carrier 1 |
| ↑ | Q9QZM3 | Cardiotrophin-like cytokine factor 1 | ↓ | Q8VD37 | SH3-containing GRB2-like protein 3-interacting protein 1 |
| ↑ | Q922S4 | cGMP-dependent 3',5'-cyclic phosphodiesterase | ↓ | Q80V94 | AP-4 complex subunit epsilon-1 |
| ↑ | Q9R087 | Glypican-6 | ↓ | Q9EP89 | Serine beta-lactamase-like protein LACTB, mitochondrial |
| ↑ | Q8C0L9 | Glycerophosphocholine phosphodiesterase GPCPD1 | ↓ | Q9ET01 | Glycogen phosphorylase, liver form |
| ↑ | Q3V300 | Kinesin-like protein KIF22 | ↓ | P08905 | Lysozyme C-2 |
| ↑ | O09164 | Extracellular superoxide dismutase [Cu-Zn] | ↓ | P49935 | Pro-cathepsin H |
| ↑ | Q9Z1B3 | 1-phosphatidylinositol 4,5-bisphosphate phosphodiesterase beta-1 | ↓ | Q61086 | Frizzled-3 |

| | | | | | |
|---|---|---|---|---|---|
| ↑ | Q8K012 | Formin-binding protein 1-like | ↓ | Q8VHK5 | Membrane protein MLC1 |
| ↑ | P36895 | Bone morphogenetic protein receptor type-1A | ↓ | Q99KK1 | Receptor expression-enhancing protein 3 |
| ↑ | Q99P69 | Kinetochore protein Nuf2 | ↓ | Q99N28 | Cell adhesion molecule 3 |
| ↑ | Q9JLC3 | Methionine-R-sulfoxide reductase B1 | ↓ | P18406 | CCN family member 1 |
| ↑ | Q9CS84 | Neurexin-1 | ↓ | Q8VHJ5 | Serine/threonine-protein kinase MARK1 |
| ↑ | O70126 | Aurora kinase B | ↓ | Q08639 | Transcription factor Dp-1 |
| ↑ | Q9ES46 | Beta-parvin | ↓ | P50285 | Dimethylaniline monooxygenase [N-oxide-forming] 1 |
| ↑ | Q8BW70 | Ubiquitin carboxyl-terminal hydrolase 38 | ↓ | P0DP99 | Short transmembrane mitochondrial protein 1 |
| ↑ | P39689 | Cyclin-dependent kinase inhibitor 1 | ↓ | Q80UZ2 | Protein SDA1 homolog |
| ↑ | P31361 | POU domain, class 3, transcription factor 3 | ↓ | Q80V91 | Probable E3 ubiquitin-protein ligase DTX3 |
| ↑ | Q80TQ5 | Pleckstrin homology domain-containing family M member 2 | ↓ | P63260 | Actin, cytoplasmic 2 |
| ↑ | Q9WTR5 | Cadherin-13 | ↓ | Q497I4 | Keratin, type I cuticular Ha5 |
| ↑ | A2RSX7 | tRNA wybutosine-synthesizing protein 5 | ↓ | Q9CXE2 | B-cell CLL/lymphoma 7 protein family member A |
| ↑ | Q6PGF3 | Mediator of RNA polymerase II transcription subunit 16 | ↓ | Q9ERE2 | Keratin, type II cuticular Hb1 |
| ↑ | Q61387 | Cytochrome c oxidase subunit 7A-related protein, mitochondrial | ↓ | Q8VD57 | Vesicle transport protein SFT2B |
| ↑ | Q07832 | Serine/threonine-protein kinase PLK1 | ↓ | Q80ZD3 | Sodium-independent sulfate anion transporter |

| | | | | | |
|---|---|---|---|---|---|
| ↑ | Q9CR75 | Tumor necrosis factor receptor superfamily member 12A | ↓ | Q9D2H2 | Adenylate kinase 7 |
| ↑ | O70458 | Oncostatin-M-specific receptor subunit beta | ↓ | Q3TSG4 | RNA demethylase ALKBH5 |
| ↑ | Q6NSQ7 | Protein LTV1 homolog | ↓ | Q9QZU9 | Ubiquitin/ISG15-conjugating enzyme E2 L6 |
| ↑ | O08811 | General transcription and DNA repair factor IIH helicase subunit XPD | ↓ | Q6PEM8 | Proton-coupled folate transporter |
| ↑ | Q8BLH7 | HIRA-interacting protein 3 | ↓ | P47806 | Zinc finger protein GLI1 |
| ↑ | O35955 | Proteasome subunit beta type-10 | ↓ | Q9DAM9 | Fibronectin type 3 and ankyrin repeat domains 1 protein |
| ↑ | Q7TPD3 | Roundabout homolog 2 | ↓ | Q62276 | Mediator of RNA polymerase II transcription subunit 22 |
| ↑ | Q8VHY0 | Chondroitin sulfate proteoglycan 4 | ↓ | Q61765 | Keratin, type I cuticular Ha1 |
| ↑ | Q8CDI7 | Coiled-coil domain-containing protein 150 | ↓ | Q9ERV1 | E3 ubiquitin-protein ligase makorin-2 |
| ↑ | Q9CZB9 | Transmembrane protein 128 | ↓ | Q8VIB3 | Type 2 lactosamine alpha-2,3-sialyltransferase |
| ↑ | Q8BIE6 | FERM domain-containing protein 4A | ↓ | Q9DB41 | Mitochondrial glutamate carrier 2 |
| ↑ | Q80ZM8 | Cardiolipin synthase (CMP-forming) | ↓ | Q64523 | Histone H2A type 2-C |
| ↑ | Q8VC74 | Cytochrome c oxidase assembly protein COX18, mitochondrial | ↓ | Q3THW5 | Histone H2A.V |
| ↑ | Q8CI43 | Myosin light chain 6B | ↓ | O70309 | Integrin beta-5 |

| | | | | | |
|---|---|---|---|---|---|
| ↑ | Q8R2U7 | Leucine-rich repeat-containing protein 42 | ↓ | Q9CRB6 | Tubulin polymerization-promoting protein family member 3 |
| ↑ | Q08274 | Dystrophia myotonica WD repeat-containing protein | ↓ | P17156 | Heat shock-related 70 kDa protein 2 |
| ↑ | Q6NVG1 | Lysophospholipid acyltransferase LPCAT4 | ↓ | Q925N0 | Sideroflexin-5 |
| ↑ | Q91X96 | Guanine nucleotide exchange factor MSS4 | ↓ | Q8BZF8 | Phosphoglucomutase-like protein 5 |
| ↑ | Q922S8 | Kinesin-like protein KIF2C | ↓ | Q9D7V9 | N-acylethanolamine-hydrolyzing acid amidase |
| ↑ | P21803 | Fibroblast growth factor receptor 2 | ↓ | P52019 | Squalene monooxygenase |
| ↑ | Q9CPV7 | Palmitoyltransferase ZDHHC6 | ↓ | P24452 | Macrophage-capping protein |
| ↑ | Q9DBR2 | Protein FAM13C | ↓ | Q99J56 | Derlin-1 |
| ↑ | P30280 | G1/S-specific cyclin-D2 | ↓ | Q9EP96 | Solute carrier organic anion transporter family member 1A4 |
| ↑ | Q8BMQ8 | Vacuolar fusion protein MON1 homolog B | ↓ | P62320 | Small nuclear ribonucleoprotein Sm D3 |
| ↑ | Q3KNJ2 | Non-homologous end-joining factor 1 | ↓ | Q60648 | Ganglioside GM2 activator |
| ↑ | Q9CWR0 | Rho guanine nucleotide exchange factor 25 | ↓ | Q9D2C7 | Bax inhibitor 1 |
| ↑ | P59328 | WD repeat and HMG-box DNA-binding protein 1 | ↓ | P16330 | 2',3'-cyclic-nucleotide 3'-phosphodiesterase |
| ↑ | Q61009 | Scavenger receptor class B member 1 | ↓ | Q9ESM6 | Glycerophosphoinositol inositolphosphodiesterase GDPD2 |
| ↑ | Q9CW79 | Golgin subfamily A member 1 | ↓ | Q80VQ0 | Aldehyde dehydrogenase family 3 member B1 |

| | | | | | |
|---|---|---|---|---|---|
| ↑ | A3KGW5 | Inactive glycosyltransferase 25 family member 3 | ↓ | Q9EPK2 | Protein XRP2 |
| ↑ | P16056 | Hepatocyte growth factor receptor | ↓ | Q8BFZ3 | Beta-actin-like protein 2 |
| ↑ | Q8CGI1 | Protein FAM193A | ↓ | P03995 | Glial fibrillary acidic protein |
| ↑ | Q80VJ2 | Steroid receptor RNA activator 1 | ↓ | Q61233 | Plastin-2 |
| ↑ | Q8BH64 | EH domain-containing protein 2 | ↓ | Q925G2 | Plasma membrane ascorbate-dependent reductase CYBRD1 |
| ↑ | Q62351 | Transferrin receptor protein 1 | ↓ | P50429 | Arylsulfatase B |
| ↑ | P52624 | Uridine phosphorylase 1 | ↓ | Q8R164 | Valacyclovir hydrolase |
| ↑ | P62071 | Ras-related protein R-Ras2 | ↓ | P11798 | Calcium/calmodulin-dependent protein kinase type II subunit alpha |
| ↑ | Q9D083 | Kinetochore protein Spc24 | ↓ | Q8K0B2 | Lysosomal cobalamin transport escort protein LMBD1 |
| ↑ | P52293 | Importin subunit alpha-1 | ↓ | Q9DB26 | Phytanoyl-CoA dioxygenase domain-containing protein 1 |
| ↑ | P11440 | Cyclin-dependent kinase 1 | ↓ | Q9EQG7 | Ectonucleotide pyrophosphatase/phosphodiesterase family member 5 |
| ↑ | P47877 | Insulin-like growth factor-binding protein 2 | ↓ | P08730 | Keratin, type I cytoskeletal 13 |
| ↑ | O08663 | Methionine aminopeptidase 2 | ↓ | Q64435 | UDP-glucuronosyltransferase 1-6 |
| ↑ | P14873 | Microtubule-associated protein 1B | ↓ | Q9CYH2 | Peroxiredoxin-like 2A |
| ↑ | Q7TN75 | Retrotransposon-derived protein PEG10 | ↓ | Q62252 | Sperm surface protein Sp17 |

| | | | | | |
|---|---|---|---|---|---|
| ↑ | P49717 | DNA replication licensing factor MCM4 | ↓ | Q9ESW8 | Pyroglutamyl-peptidase 1 |
| ↑ | P49718 | DNA replication licensing factor MCM5 | ↓ | Q9D3P8 | Plasminogen receptor (KT) |
| ↑ | P24547 | Inosine-5'-monophosphate dehydrogenase 2 | ↓ | Q91Z53 | Glyoxylate reductase/hydroxypyruvate reductase |
| ↑ | P13864 | DNA (cytosine-5)-methyltransferase 1 | ↓ | Q9Z0J0 | NPC intracellular cholesterol transporter 2 |
| ↑ | Q61881 | DNA replication licensing factor MCM7 | ↓ | Q64475 | Histone H2B type 1-B |
| ↑ | P14069 | Protein S100-A6 | ↓ | Q9QXE0 | 2-hydroxyacyl-CoA lyase 1 |
| ↑ | Q8BTG7 | Protein NDRG4 | ↓ | Q8C0M9 | Isoaspartyl peptidase/L-asparaginase |
| ↑ | P97311 | DNA replication licensing factor MCM6 | ↓ | P10922 | Histone H1.0 |
| ↑ | P01831 | Thy-1 membrane glycoprotein | ↓ | Q3U5Q7 | UMP-CMP kinase 2, mitochondrial |
| ↑ | P11157 | Ribonucleoside-diphosphate reductase subunit M2 | ↓ | Q9D5Y1 | Coiled-coil domain-containing protein 39 |
| ↑ | Q3U962 | Collagen alpha-2(V) chain | ↓ | Q00915 | Retinol-binding protein 1 |
| ↑ | E9Q394 | A-kinase anchor protein 13 | ↓ | Q8BS45 | Intraflagellar transport protein 56 |
| ↑ | Q61704 | Inter-alpha-trypsin inhibitor heavy chain H3 | ↓ | Q91XE8 | Transmembrane protein 205 |
| ↑ | Q3UZ39 | Leucine-rich repeat flightless-interacting protein 1 | ↓ | P70665 | Sialate O-acetylesterase |
| ↑ | O09110 | Dual specificity mitogen-activated protein kinase 3 | ↓ | O54782 | Epididymis-specific alpha-mannosidase |

| | | | | | |
|---|---|---|---|---|---|
| ↑ | O70131 | Ninjurin-1 | ↓ | Q9DBB8 | Trans-1,2-dihydrobenzene-1,2-diol dehydrogenase |
| ↑ | Q9Z127 | Large neutral amino acids transporter small subunit 1 | ↓ | P84104 | Serine/arginine-rich splicing factor 3 |
| ↑ | P25206 | DNA replication licensing factor MCM3 | ↓ | P83510 | Traf2 and NCK-interacting protein kinase |
| ↑ | P06837 | Neuromodulin | ↓ | P97821 | Dipeptidyl peptidase 1 |
| ↑ | Q01320 | DNA topoisomerase 2-alpha | ↓ | P62717 | 60S ribosomal protein L18a |
| ↑ | Q8VDW0 | ATP-dependent RNA helicase DDX39A | ↓ | P15864 | Histone H1.2 |
| ↑ | Q9R0E1 | Multifunctional procollagen lysine hydroxylase and glycosyltransferase LH3 | ↓ | Q99L20 | Glutathione S-transferase theta-3 |
| ↑ | Q8K2Z4 | Condensin complex subunit 1 | ↓ | Q8BFS6 | Serine/threonine-protein phosphatase CPPED1 |
| ↑ | Q62219 | Transforming growth factor beta-1-induced transcript 1 protein | ↓ | Q9ET22 | Dipeptidyl peptidase 2 |
| ↑ | Q64152 | Transcription factor BTF3 | ↓ | Q8BTJ4 | Bis(5'-adenosyl)-triphosphatase enpp4 |
| ↑ | Q60715 | Prolyl 4-hydroxylase subunit alpha-1 | ↓ | Q3U0V2 | Tumor necrosis factor receptor type 1-associated DEATH domain protein |
| ↑ | P13439 | Uridine 5'-monophosphate synthase | ↓ | Q9CQY6 | Ubiquinol-cytochrome-c reductase complex assembly factor 2 |
| ↑ | Q64337 | Sequestosome-1 | ↓ | P21956 | Lactadherin |
| ↑ | O89086 | RNA-binding protein 3 | | | |
| ↑ | Q91Z31 | Polypyrimidine tract-binding protein 2 | | | |

| | | |
|---|---|---|
| ↑ | Q9WTI7 | Unconventional myosin-Ic |
| ↑ | Q8C156 | Condensin complex subunit 2 |
| ↑ | Q91VJ2 | Caveolae-associated protein 3 |
| ↑ | P97825 | Jupiter microtubule associated homolog 1 |
| ↑ | Q61730 | Interleukin-1 receptor accessory protein |
| ↑ | P55821 | Stathmin-2 |
| ↑ | Q3UA16 | Kinetochore protein Spc25 |

**Table S2. List of up- and downregulated proteins in *Mlc1*-null astrocytes compared to wild-type astrocytes.** 134 up- and 125 downregulated proteins based on a fold change > 0.06 or < -0.06 with an adj. $P<0.05$.
